# Inhibition of Endothelial Lipase by MEDI5884 Normalizes Phosphatidylinositol Levels in Coronary Artery Disease Patients

**DOI:** 10.1101/2024.05.30.596497

**Authors:** Anton I. Rosenbaum, Yue Huang, Ruipeng Mu, Kristina Kovacina, ChaoYu Denise Jin, B. Timothy Hummer, Meina Liang, Joseph S. Grimsby

## Abstract

**Background:** Endothelial lipase (EL) promotes high-density lipoproteins (HDL) phospholipid degradation, increases catabolism of HDL and is an attractive target for the potential treatment for cardiovascular disease. Inhibition of EL using a monoclonal neutralizing antibody, MEDI5884, demonstrated increased quantity and function of HDL. Determinants of anti-atherosclerotic function of HDL comprise the interplay of various components of HDL structure-activity relationship: size, shape and composition (lipid and protein). Previous studies have shown that single doses of MEDI5884 administered to healthy nonhuman primates (NHPs) and healthy subjects resulted in a dose-dependent increase in plasma phospholipids (PL) and that plasma PI levels in placebo treated healthy subjects are significantly increased relative to CAD subjects participating in clinical trials NCT03001297 and NCT03351738, respectively.

**Methods:** Herein, we characterized using LC-MS/MS the plasma lipidome of NHPs, heathy subjects and subjects with coronary artery disease (CAD) following MEDI5884 administration.

**Results:** MEDI5884 treated NHPs resulted in a prominent increase in phosphatidylinositols (PI) and cholesteryl esters (CE). Treatment with MEDI5884 restores near-normal levels of PI in CAD patients. PI increases in both healthy subjects and CAD patients were dose-dependent, correlated with exposure and saturated at approximately 200 mg MEDI5884 subcutaneous (SC) dose in CAD patients. Comparison of pharmacodynamic (PD) effects of repeat SC 200 mg doses of MEDI5884 in CAD patients revealed greater and more rapid increases in PI levels compared to HDL-C and HDL phospholipid (HDL-PL). The increase in PI species was inversely correlated with decreases in free EL mass levels.

**Conclusions:** PI has previously been shown to possess anti-atherosclerotic properties and led to increases in HDL cholesterol (HDL-C) and reverse cholesterol transport (RCT). The mechanism by which CE levels increase as the result of MEDI5884 administration can be attributed to the observed increase in both substrates of the lecithin-cholesterol acyltransferase (LCAT) reaction: phosphatidylcholine/phosphatidylethanolamine (PC/PE) and cholesterol as the consequence of EL inhibition. Further characterization of the underlying biological mechanisms responsible for the decrease of the PI biomarker in CAD patient population relative to healthy subjects as well as in conjunction with pharmacological intervention by MEDI5884 may reveal more information on this clinically-relevant biomarker and potential role in CAD.

## Introduction

HDL removes cholesterol from macrophages and peripheral tissues and delivers it to the liver directly via scavenger receptor class B type 1 (SRB1) receptor or via low-density lipoproteins LDL and the LDL receptor for recycling or elimination via bile [1].

Endothelial lipase plays a critical role in maintaining the catabolism and homeostasis of HDL by hydrolyzing phospholipids and promoting the degradation of HDL particles and subsequent elimination in urine. MEDI5884 is a neutralizing monoclonal antibody against endothelial lipase being studied for the treatment of coronary artery disease (CAD)[2, 3]. Since the failure of cholesteryl ester transfer protein (CETP) inhibitors to bring benefit to cardiovascular disease patients as it relates to augmenting HDL-C levels one has to carefully consider understanding the functionality of HDL that goes beyond cholesterol content ([4–7]). An interplay of structure activity relationship exists for HDL where changes in its composition (proteome, lipidome) elicit changes in its structure, shape, size and anti-atherosclerotic properties ([8]).

Previous studies have shown that single doses of MEDI5884 administered to healthy monkeys resulted in a dose-dependent increase in HDL, plasma phospholipids (PL) and increased efflux. Further studies in HV and CAD patients revealed increases in HDL-C, HDL-PL and efflux [2, 9]. To elucidate the phospholipid changes resulting from the administration of MEDI5884 we undertook a mass spectrometry-based lipidomic assessment which revealed a dose-dependent increase in phosphatidylinositols upon administration of MEDI5884.

## Results

### Effects in Cynomolgus Monkeys

Lipidomic analyses revealed that PI species were substantially increased compared to other phospholipids (Figure 1) in plasma from cynomolgus monkeys upon single dose administration of MEDI5884 (30 mg/kg). Hierarchical clustering diagram revealed that PI species cluster together and increase in their levels much more drastically compared to other lipid species in a time-dependent fashion. Furthermore, this finding was confirmed using principle component analysis (PCA) which identified several sets of lipids clusters: PI and cholesterol esters (Figure 2).

**Figure 1.**
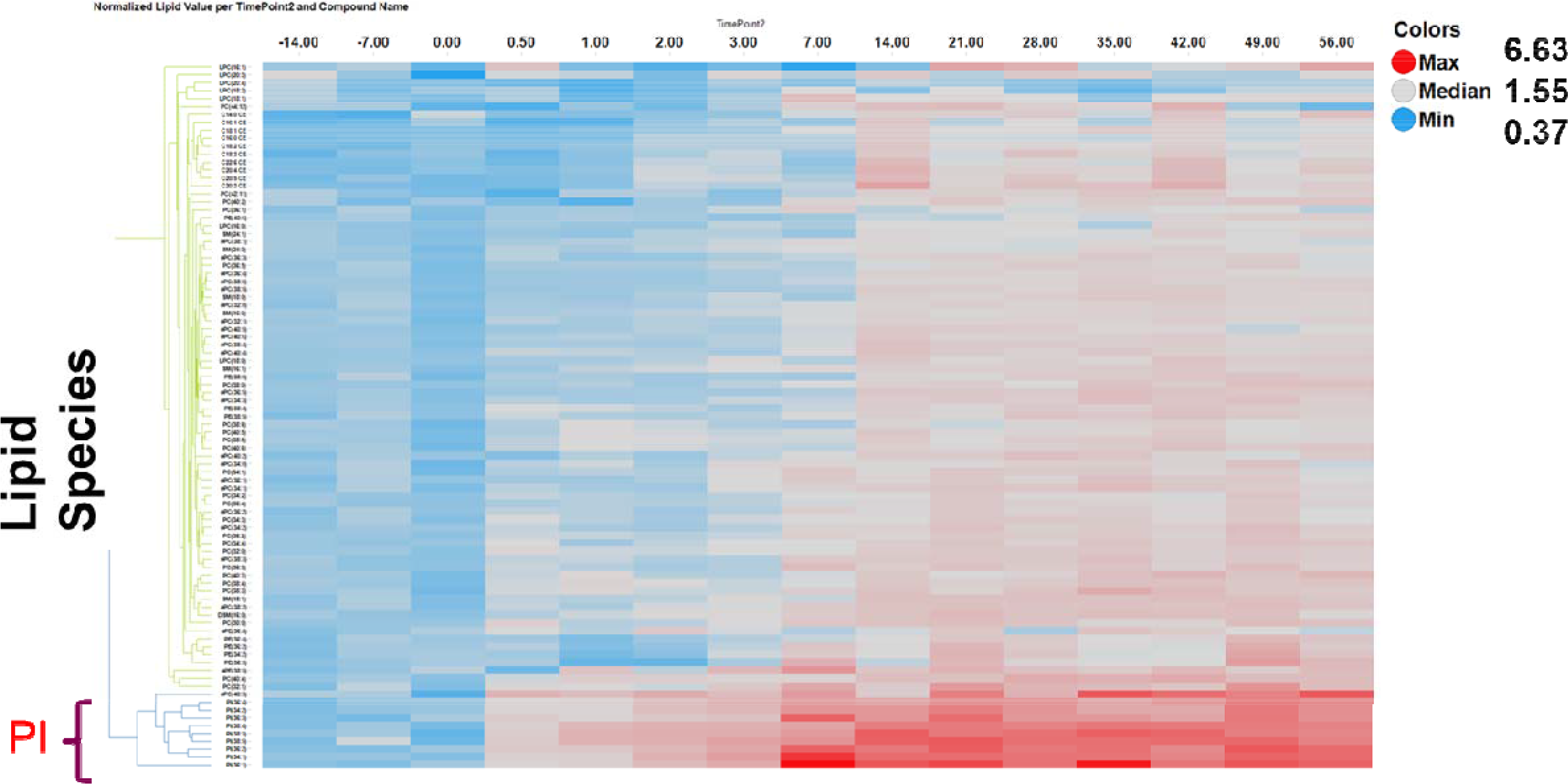
Hierarchical clustering analysis of cyno plasma phospholipidomic dataset after a single SC dose of MEDI5884 (30 mg/kg).

**Figure 2.**
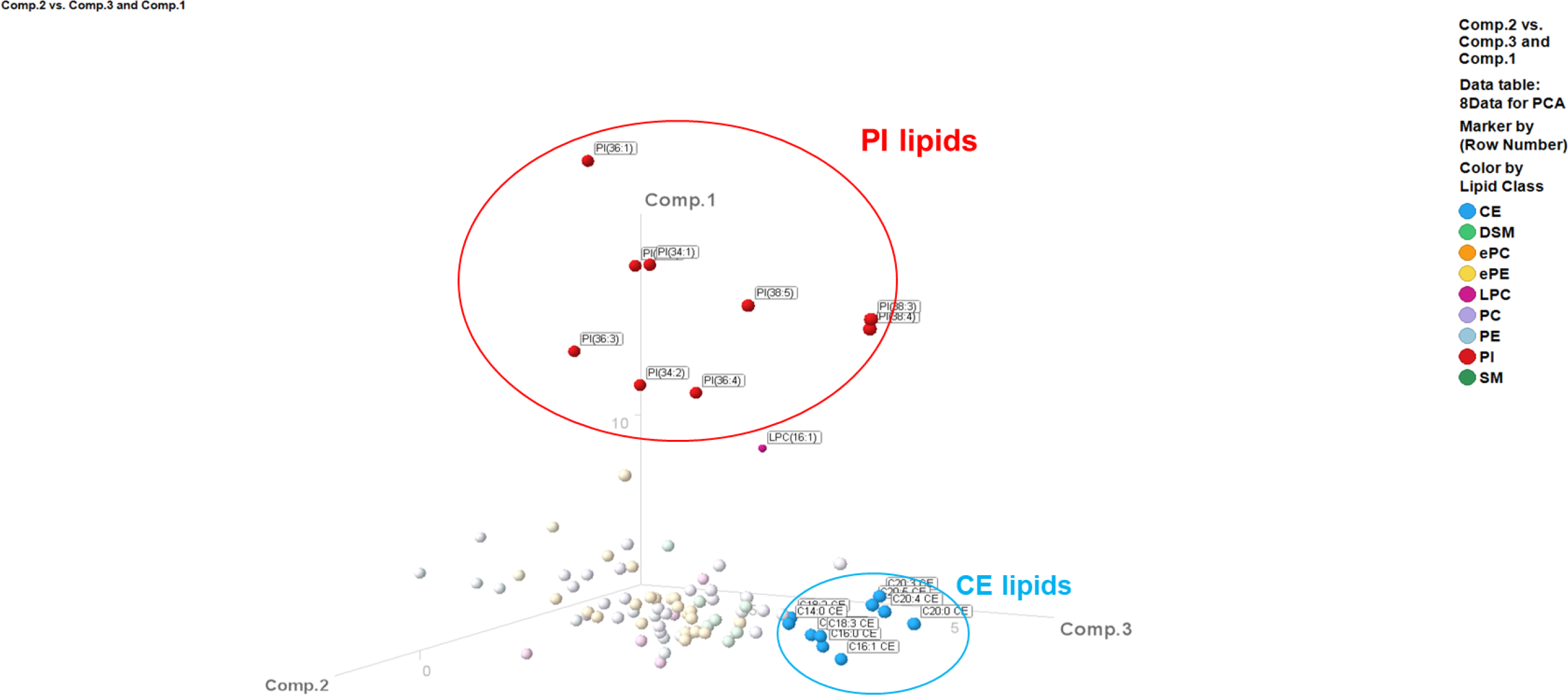
Principal component analysis (PCA) of phospholipidomic data from cynomolgus monkey plasma upon adminstration of MEDI5884 at 0.5, 6, and 30 mg/kg (all timpoints) identifies several lipid clusters: PI and CE lipids.

Cholesteryl esters are products of the lecithin–cholesterol acyltransferase (LCAT) (EC2.3.1.43) reaction where an *sn2* acyl chain derived from phosphatidylcholine (PC) or phosphatidylethanolamine (PE) is transesterified to cholesterol [10]. The products of LCAT reaction are LysoPC and cholesteryl ester. Our lipidomic data revealed that *both* substrates of the LCAT reaction: PC/PE and cholesterol increased as the result of MEDI5884 treatment resulting in increased cholesteryl ester levels (Supplemental Figure 1). Possible explanations include activation of LCAT by a >30% increase in ApoA1, enrichment of acidic phospholipid such as PI, increased recruitment of LCAT to HDL and/or increased synthesis of LCAT[11]. PI and CE lipids correlated well with concomitant HDL-C increases as shown in Supplemental Figures 2 and 3, respectively.

**Figure 3.**
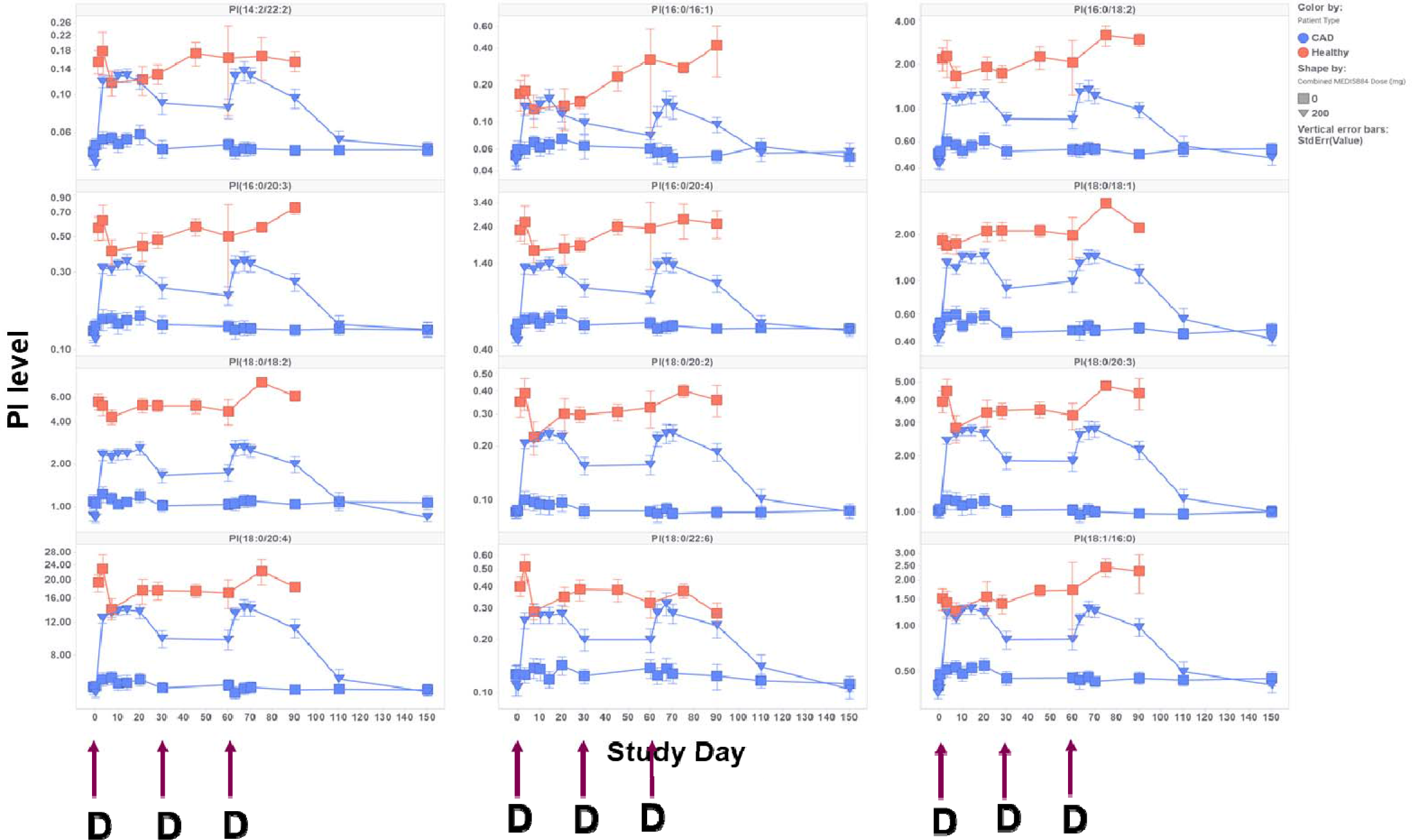
200 mg SC monthly dose of MEDI5884 in CAD patients increased PI to levels observed in healthy subjects.

### Effects in HV Subjects

Following a single subcutaneous dose of MEDI5884 in study NCT03001297, increases in the majority of plasma PI species were observed at all dose levels of MEDI5884 relative to the placebo (Supplemental Figure 4). The duration of the increases appeared to correlate with MEDI5884 exposure in a dose-dependent manner; subjects who received MEDI5884 600 mg showed the most sustained response post Day 28, which lasted until the end of the study (90 days) for some PI species. For the majority of PI species, the maximal increase in plasma PI levels occurred on Day 21 and/or Day 28 and appeared to reach a plateau in the MEDI5884 300 mg dose group where there was a minimal difference in change compared with the MEDI5884 600 mg dose group consistent with maximum complete neutralization of circulating EL.

### Effects in CAD Patients

As illustrated in Supplemental Figure 5 following 3 monthly SC doses of MEDI5884 in study NCT03351738, most plasma PI species dose-dependently increased relative to placebo. The duration of increases in plasma PI appeared to correlate with MEDI5884 exposure. For most PI species, increases in PI levels reached saturation at the MEDI5884 350 mg dose level, and the MEDI5884 500 mg dose level did not result in further PI increases relative to baseline. However, on Day 91 at the MEDI5884 200 mg dose level, PI increases approached saturation levels observed in higher dose cohorts.

Previous studies have shown that PI lipids are substantially lower in CAD patients compared to HV subjects[12]. We have endeavored to examine the effects of MEDI5884 on PI species in CAD patients relative to HV subjects. As is shown in **Figure 3**, a 200 mg SC monthly dose of MEDI5884 in CAD patients increased PI to levels observed in healthy subjects thus normalizing their PI levels.

### Effects of MEDI5884 on Free Endothelial Lipase Levels in HV Subjects and CAD Patients

Following administration of MEDI5884, dose-dependent suppression of EL levels was observed (Supplemental Figures 6, 7 and 8). Complete suppression of EL levels was observed for up to 45 days post dosing for the MEDI5884 300 and 600 mg dose groups for HV. In CAD subjects, after 3 monthly SC MEDI5884 doses, dose-dependent suppression of EL levelswas observed. EL levelsdecreased >85% from baseline through Day 91 at MEDI5884 350 and 500 mg doses. EL levels decreased >60% from baseline through Day 91 at MEDI5884 200 mg. Initially, the 100 mg dose suppressed free EL nearly completely, while the 50 mg dose achieved approximately 80% suppression. Subsequently, EL suppression reversed consistently as MEDI5884 exposure decreased, in a dose-dependent manner.Greater and more rapid increases in PI levels compared to HDL-C and HDL-PL were observed, consistent with exposure to MEDI5884 [9] and with concomitant decreases in free EL mass levels (**Figure 4**).

**Figure 4.**
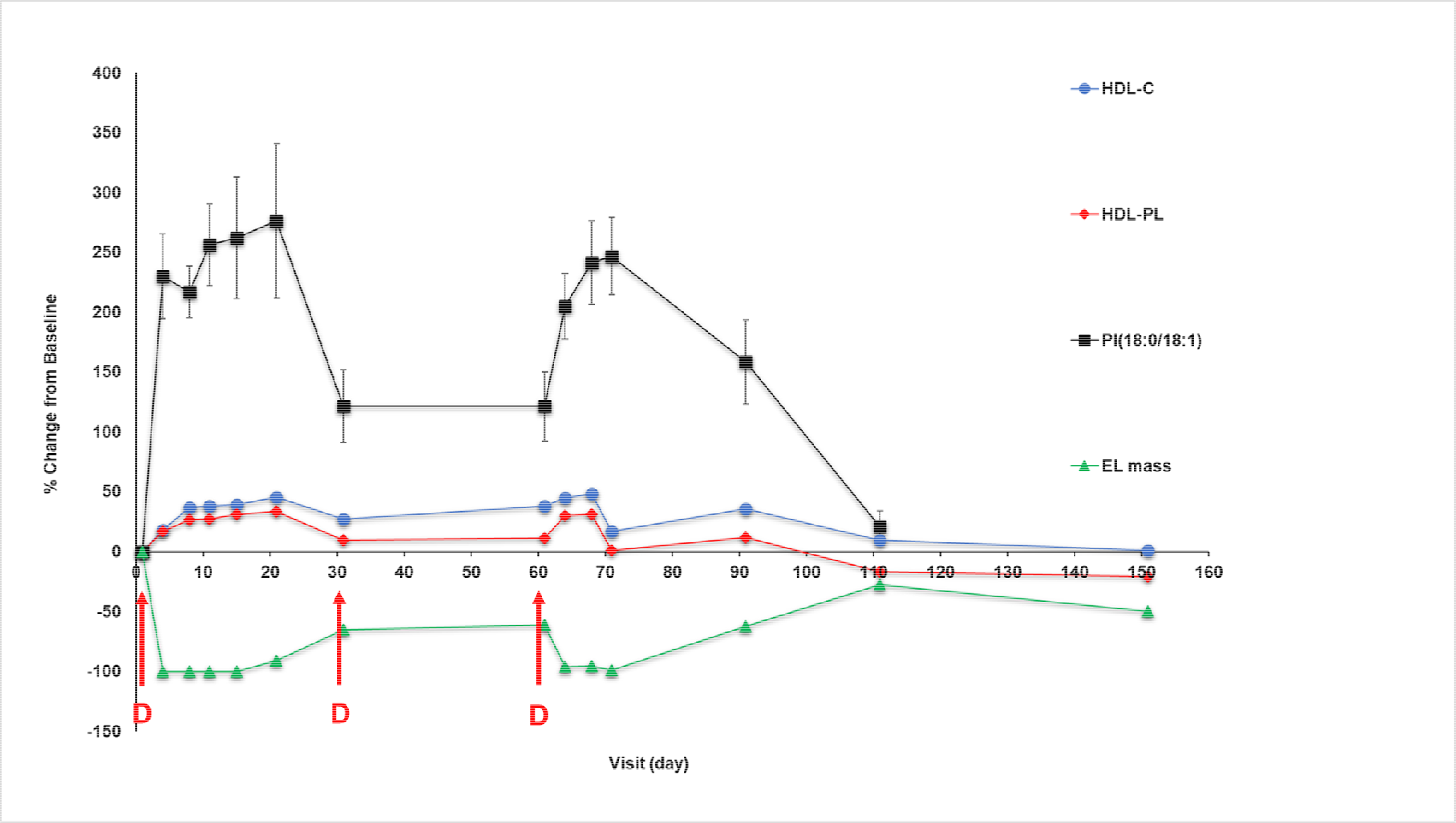
Comparison of pharmacodynamic effects of 3 repeat monthly SC 200 mg doses of MEDI5884 (denoted with arrows) in CAD patients. Pharmacodynamic biomarkers were HDL-C, HDL-PL, PI (18:0/18:1) and free EL mass, which were evaluated after the first and third MEDI5884 administrations.

## Discussion

Early studies have shown that injection of PI into rabbits promotes cholesterol elimination and affects surface potential/charge of various lipoprotein particles, including HDL [13]. Further studies have shown that PI promotes cholesterol efflux from J774 cells and yet again injection of PI into rabbits promotes excretion of tritiated cholesterol into bile indicating increased reversed cholesterol transport. Additionally, PI enriched HDL increases cholesterol update into HPG2 cells compared to PC-rich HDL [14].

Previous studies have indicated that changes in lipid composition of HDL can affect its bioactivity [15]. PI along with other negatively charged phospholipids may impact the net surface charge of HDL thereby modulating charge-dependent interactions with lipases, lipid transfer proteins, extracellular matrix, and other protein components (PMID: 23543772). PI has been shown to alter the physical state of phosphatidylcholine synthetic membranes and promote fluid phase formation and packing disorder [16].

Additional studies have demonstrated an intermolecular interaction of phosphatidylinositol with the lipid raft molecules sphingomyelin and cholesterol [17]. These data indicate that PI can affect cholesterol chemical potential thus promoting its efflux. Cholesterol chemical potential is an important variable in affecting its transport and trafficking [18]. Additionally, administration of PI has been shown to increase HDL-C levels in humans [19]. Furthermore, an association between low HDL-PI in subjects with CAD status and high HDL-C have been reported previously [20]. Additionally, PI has been shown to be decreased in acute-phase HDL (APHDL) obtained 34–38 h after surgery from patients who underwent bypass surgical procedures [21, 22].

Interestingly, Stamler et al [13] have shown that PI inhibits esterification by LCAT and reduces CE levels. This differs from our observation in cynomolgus monkeys where we have observed increased CE levels as the result of MEDI5884 administration and concomitant PI level increases. This difference could be due to the fact we also observed PC/PE level increases as the result of MEDI5884 administration. As mentioned above PC/PE are substrates for the LCAT reaction, and PI is not. One possible explanation is that in experiments conducted by Stamler et al. [13] reduction in PC as one of the substrates required for CE formation by LCAT due to administration of PI led to the reduced esterification rate.This could be especially true since LCAT activity was measured using radioactive free cholesterol, not radioactive PC/PE. Additional explanations include the potential species differences and pleiotropic effects of MEDI5884 administration.

The literature taken together indicates that PI is an important phospholipid that can modulate lipoprotein particle charge, chemical potential of membranes and affect cholesterol transport in vivo promoting its elimination. Previous studies [2, 9] have shown that MEDI5884 treatment resulted in not only increased HDL-C quantity but also improved quality as demonstrated by increased cholesterol efflux capacity. In conjunction with the finding that MEDI5884 promotes cholesterol efflux we can hypothesize that changes in plasma lipidome, in particular elevation of PI levels, as the result of administration of MEDI5884 alters the structure activity relationship of HDL particles to promote cholesterol efflux from macrophages.

In conclusion, treatment with MEDI5884, an endothelial lipase neutralizing monoclonal antibody being developed for the treatment of coronary artery disease by increasing HDL quantity and function, restores near-normal levels PI in CAD patients on intensive statin therapy in a dose-dependent manner. Further characterization of the underlying biological mechanisms responsible for the decrease of the PI biomarker in CAD patient population relative to healthy subjects as well as in conjunction with pharmacological intervention by MEDI5884 may reveal more information on this clinically-relevant biomarker and elucidate how changes in lipidome can affect cholesterol efflux capacity of HDL particles. Moreover, additional studies establishing that increasing PI in CAD patients impacts HDL function and clinical outcome would need to be conducted.

## Materials and Methods

Groups of 3 male cynomolgus macaques were dosed by single subcutaneous injection with 0.5 mg/kg, 6 mg/kg, and 30 mg/kg of MEDI5884 (S6F1-4P), and 6 mg/kg of S1- IgG1. Plasma samples were collected 14 and 7 days prior to dosing as well as shortly prior to dosing and approximately 0.5, 1, 2, 3, 7, 14, 21, 28, 35, 42, 49, 56 days after dose. Further details on study design have been published previously[2].

### Targeted Lipidomic Analysis for Monkey Plasma Samples

An automated electrospray ionization-tandem mass spectrometry approach was used, and data acquisition and analysis were carried out largely as described previously (Devaiah et al., 2006; Bartz et al., 2007) with some modifications as described in supplemental information.

### PI quantification for clinical analyses was performed as described previously ()

Quantification results of the levels of various PI species in healthy volunteers’ samples were based on a total of 214 samples from study NCT03001297 [2] and a total of 978 plasma samples from Coronary Artery Disease (CAD) subjects participating in clinical trial NCT03351738 [9].

### Free Endothelial Lipase Assay

Sandwich immunoassay has been developed for the detection human EL utilizing ECL readout. A MSD plate (MesoScale Discovery) was coated with 5 µg/mL of MEDI5884, at 50 µl/well, incubated at 4°C on a flat surface and subsequently washed 3x with 300 µl/well of ELISA Wash Buffer. The assay plate(s) were blocked with 150 µl/well of Block Buffer (IBB) for ≥60min on an orbital plate shaker at room temperature (RT). Reference standards (RS), quality control (QC) and negative control (NC), prepared in IBB, and test samples were added to the plates at 35 µl/well. Samples were tested using previously determined minimally required dilution (MRD, 2 for plasma samples). The assay plate(s) were incubated at RT on a plate shaker with gentle shaking for 60 minutes ± 10 minutes. Unbound analyte was removed by washing the plate(s) with ELISA wash buffer. To detect the captured analyte, 1 µg/ml of biotinylated anti-huEL monoclonal antibody (OriGene) was added at 35 µl/well and incubated for additional 60 minutes ± 10 minutes at RT on a plate shaker. Unbound detection antibody was removed by washing the plate(s). Streptavidin Sulfo-TAG was added at 35 µl/well and incubated for additional 60 minutes ± 10 minutes at RT on a plate shaker with gentle shaking. Plate(s) were washed again and Read Buffer added at 150 µl/well. Signal was read on a MSD sector imager instrument within 20 minutes. The ECL values for each plate were collected using the MSD Sector Imager. The ECL values for the reference standard curves for each plate were plotted with Softmax Pro GxP v6.4 software (Molecular Devices, Sunnyvale, CA) using a 1/y^2^-weighted 4-Parameter Logistic (4-PL) model of curve fitting. The concentrations of unknown samples were interpolated from the respective standard curves. The Softmax-derived data was then imported into Microsoft Excel Software and Spotfire (TIBCO® Spotfire® Analyst 7.9.2 HF-011 Build version 7.9.2.0.12) to generate data reports and graphs.

HDL-C and HDL-PL were analyzed as described previously [2]

## Acknowledgment

The authors are or were employees of AstraZeneca at the time this work was conducted and may hold stock ownership and/or stock options or interests in the company. This study was funded by AstraZeneca.

## Supplemental Figures

**Supplemental Figure 1.**
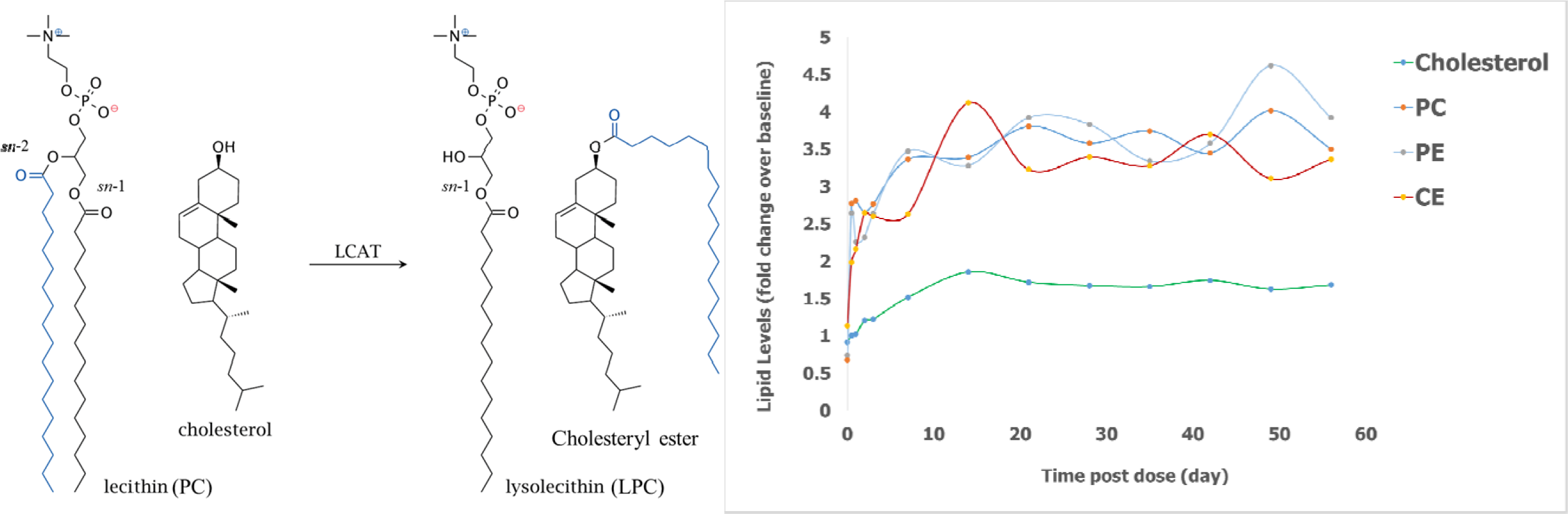
A. LCAT Reaction. B. Increase in CE as well as LCAT substrates Cholesterol, PE and PC after single SC dose of MEDI5884 (30 mg/kg) in cynomolgus monkeys.

**Supplemental Figure 2.**
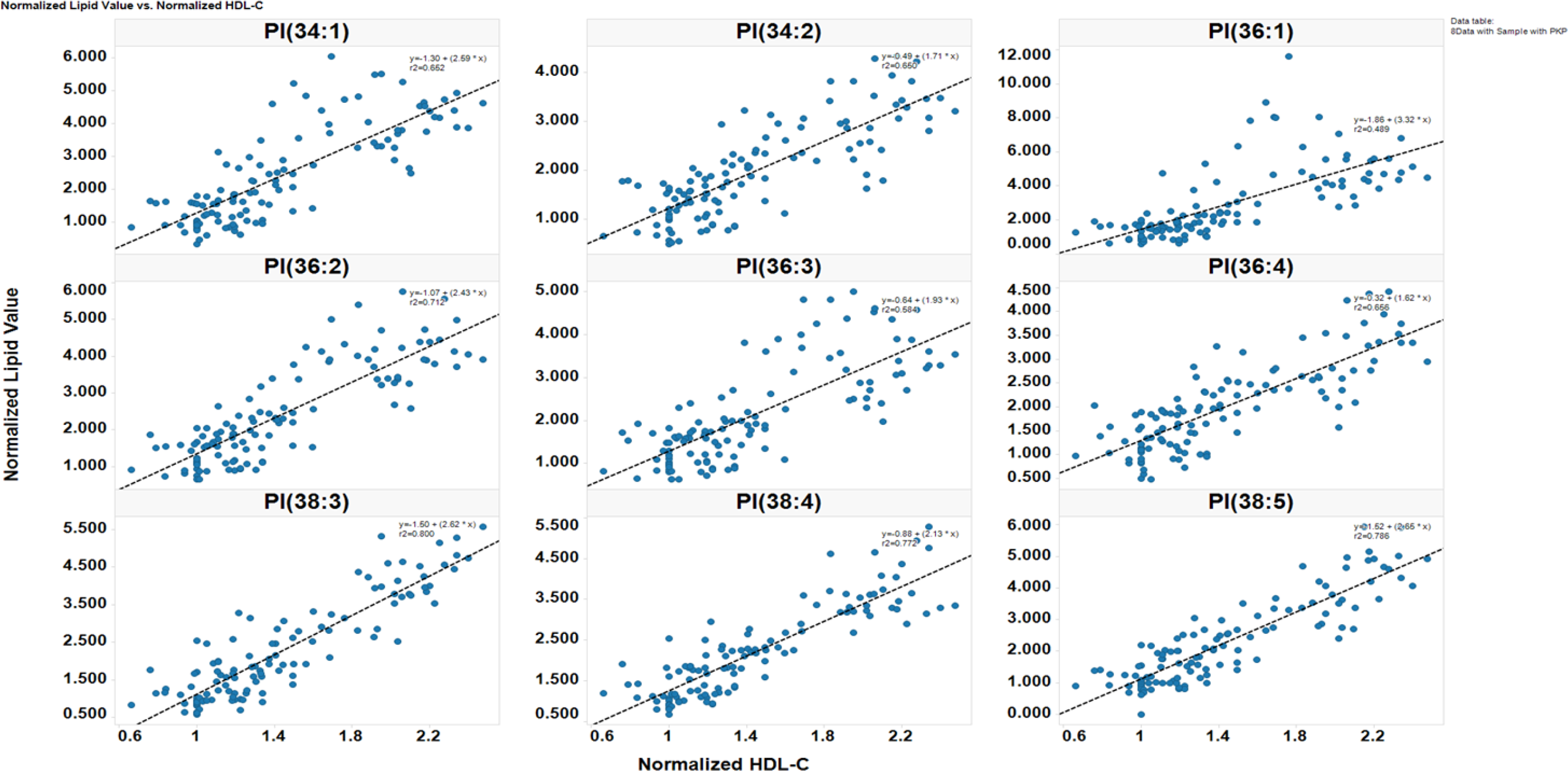
Correlation analysis of normalized PI Levels vs. normalized HDL-C Levels from cynomolgus monkeys upon administration of MEDI5884 (all dose levels and all timepoints). Analysis was performed in Spotfire.

**Supplemental Figure 3.**
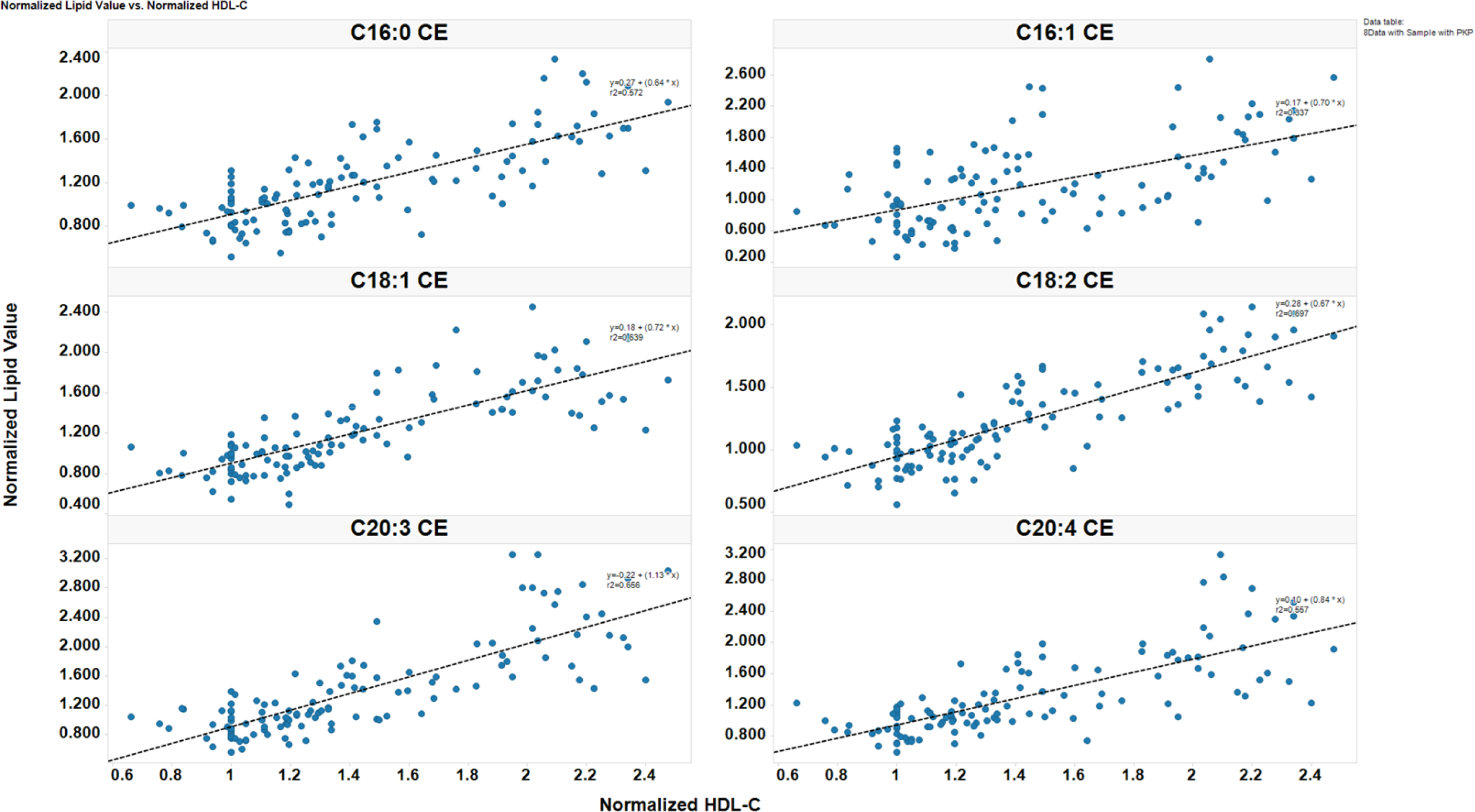
Correlation analysis of normalized cholesteryl esters levels vs. normalized HDL-C levels from cynomolgus monkeys upon administration of MEDI5884 (all dose levels and all timepoints). Analysis was performed in Spotfire.

**Supplemental Figure 4.**
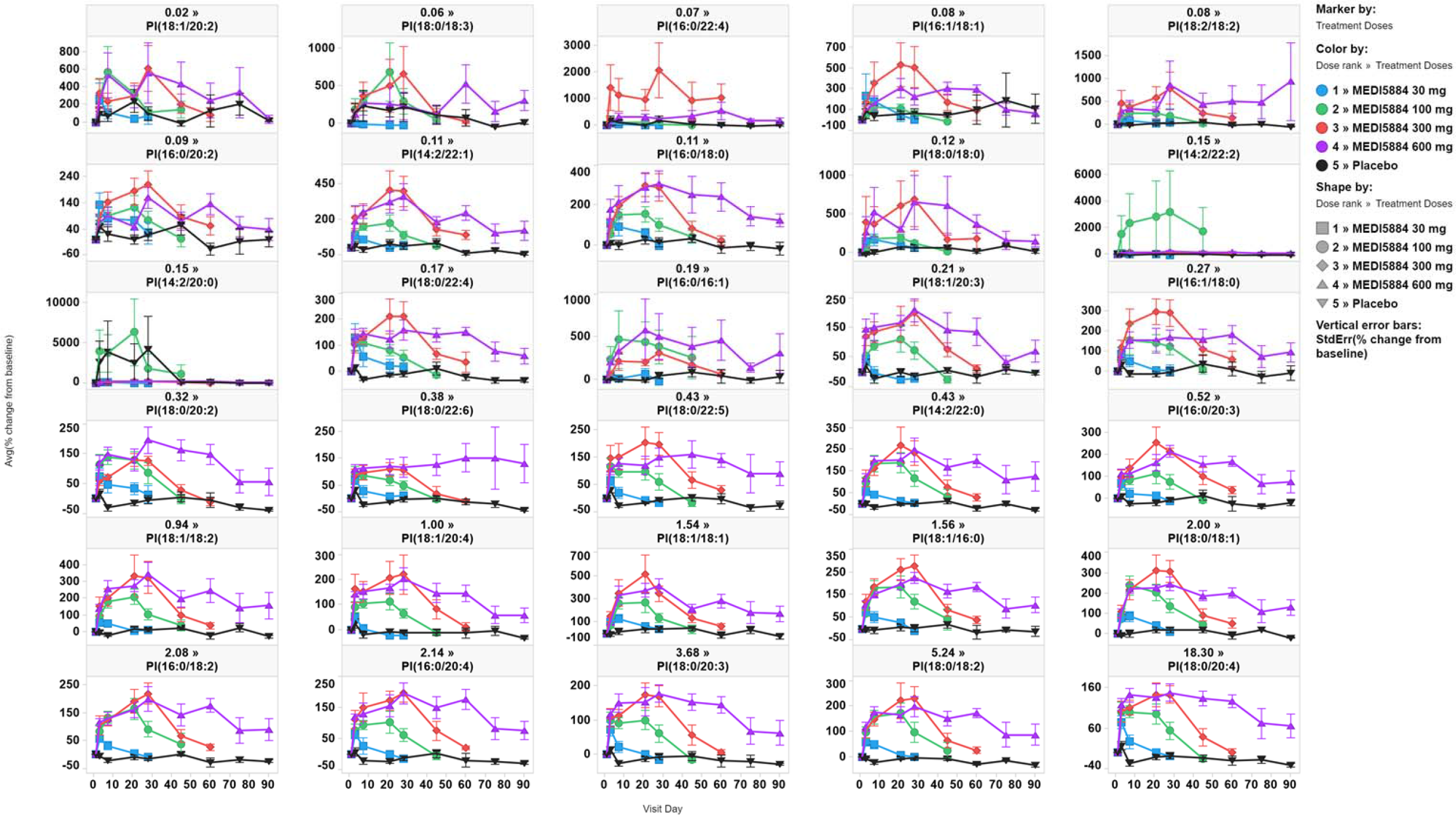
Plasma PI Area % Change from Baseline Over Time (US Population, As Treated Population) from the NCT03001297 study in HV following a single SC dose of MEDI5884. Avg = average; SEM = standard error of the mean; US = United States. Trelis legend includes a number above the PI species name, which is the average area ratio for each PI species for all timepoints for the placebo group. Error bars represent the SEM.

**Supplemental Figure 5.**
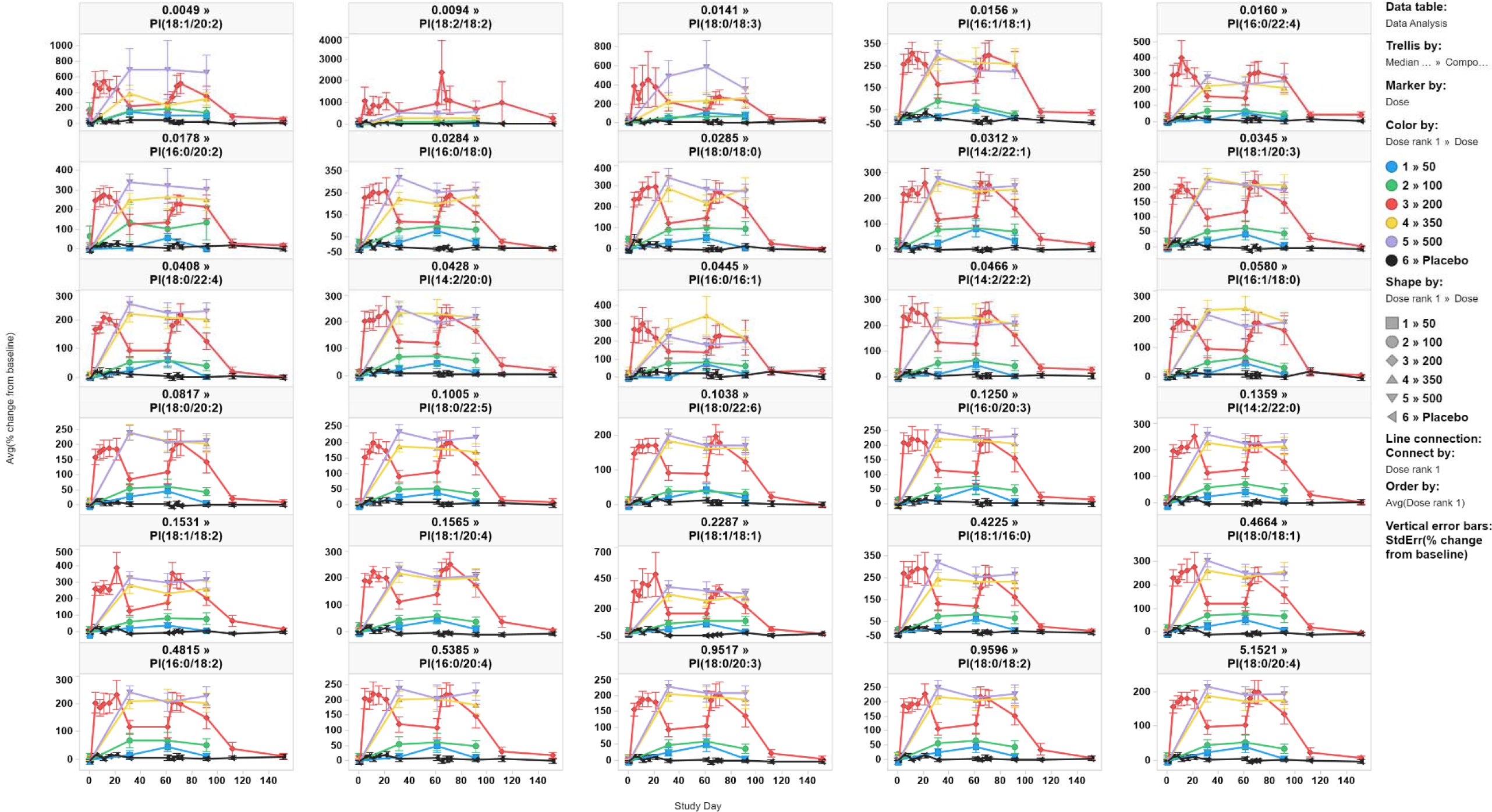
Plasma PI Area Ratio % Change from Baseline over Time for All MEDI5884 Dose Levels and Placebo (As-treated Population) from the NCT03351738 study in CAD patients. Avg = average; StdErr = standard error of the mean. Trelis legend includes a number above the PI species name, which is the average area ratio for each PI species at all time points for the placebo group. Error bars represent the StdErr. Day 1 was used as baseline. Screening visit was imputed as Day 0.MEDI5884 was administered as 3 monthly SC doses.

**Supplemental Figure 6.**
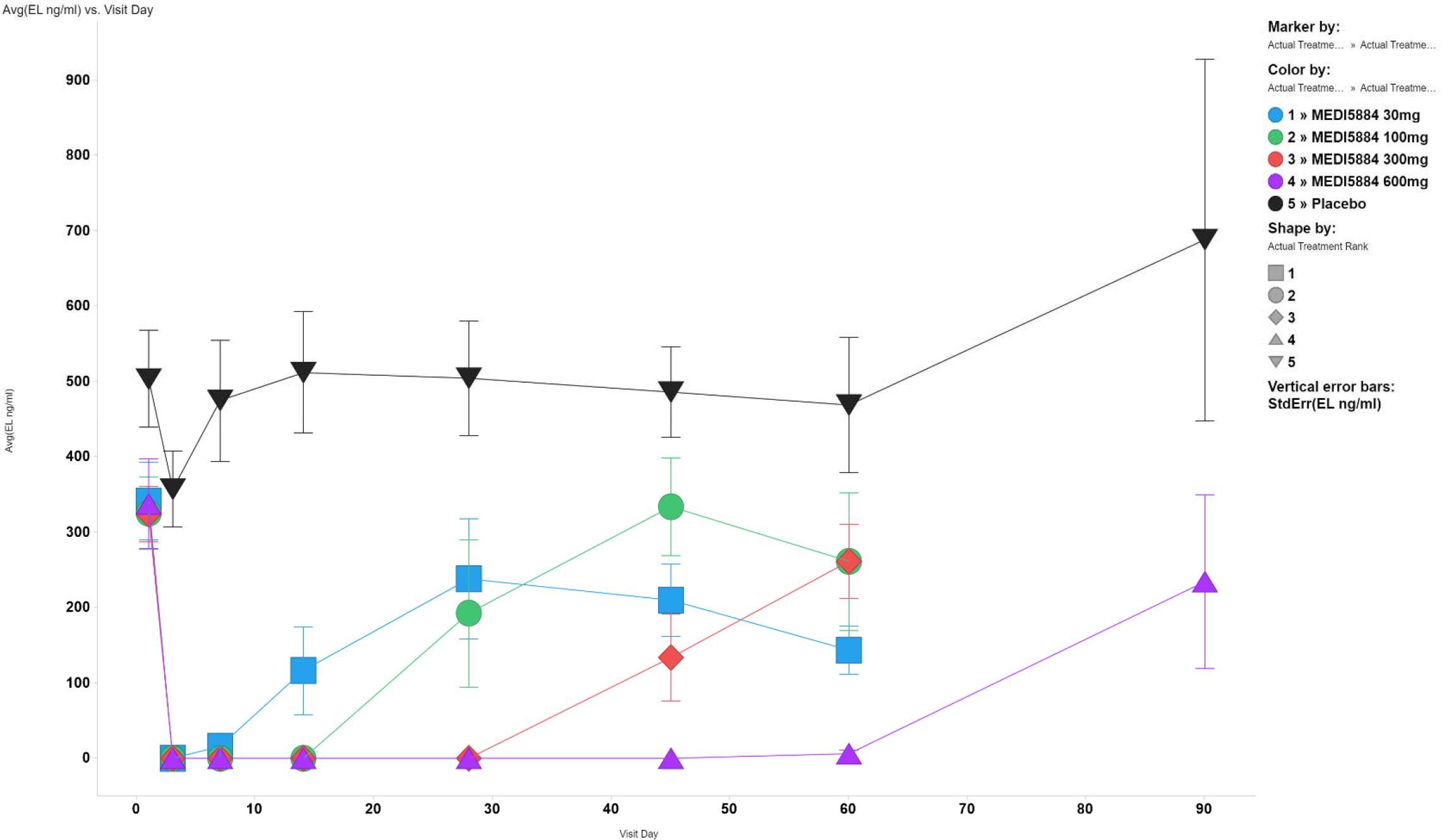
Dose-dependent suppression of EL levels in HV (As Treated Population) participating in study NCT03001297 upon administration of a single SC dose of MEDI5884. Avg = average; error bars represent SEM = standard error of the mean.

**Supplemental Figure 7.**
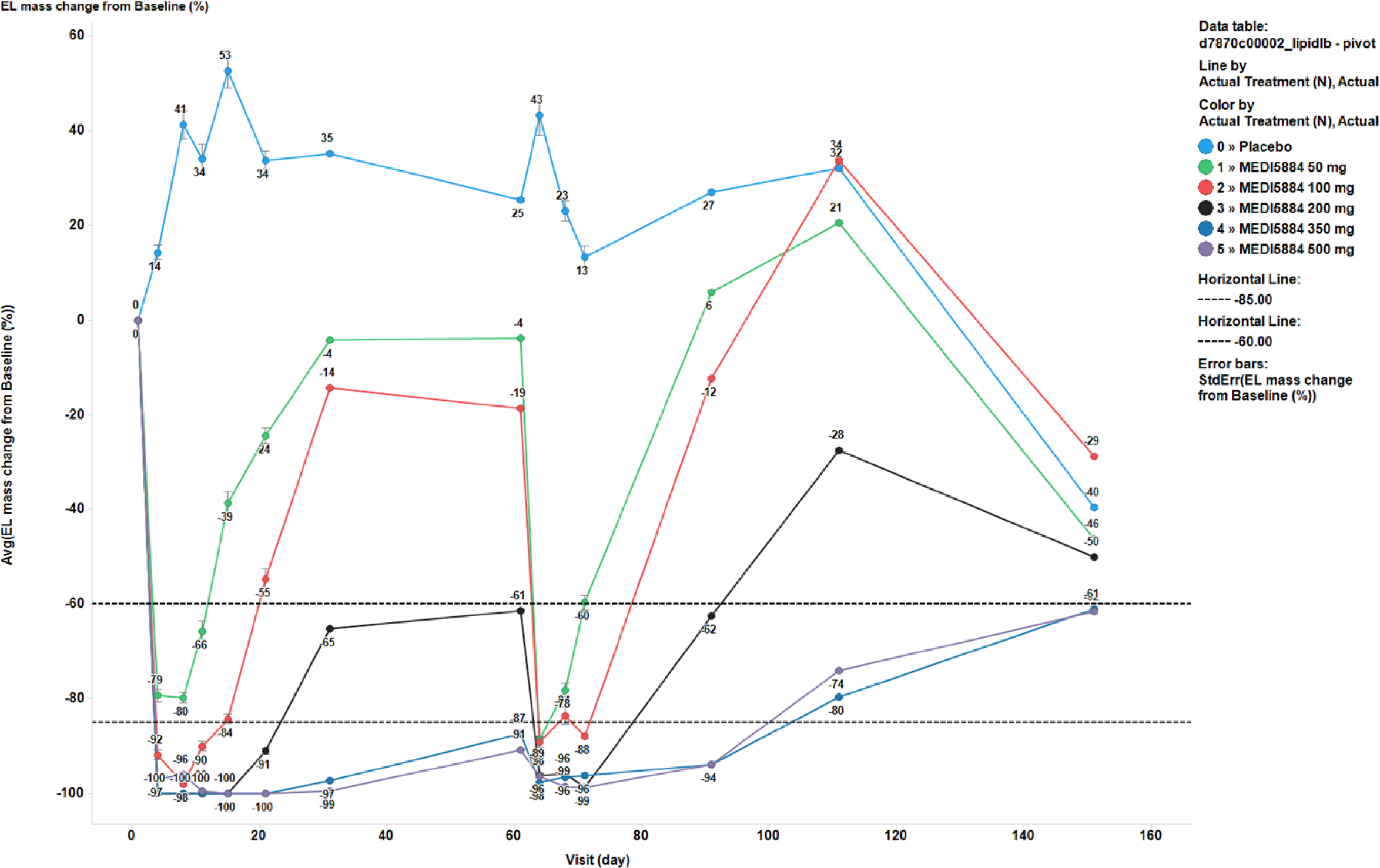
Dose-dependent suppression of EL levels (change from baseline) in CAD patients participating in study NCT03351738 upon administration of 3 monthly SC doses of MEDI5884. Avg = average; error bars represent SEM = standard error of the mean.MEDI5884 was administered as 3 monthly SC doses.

**Supplemental Figure 8.**
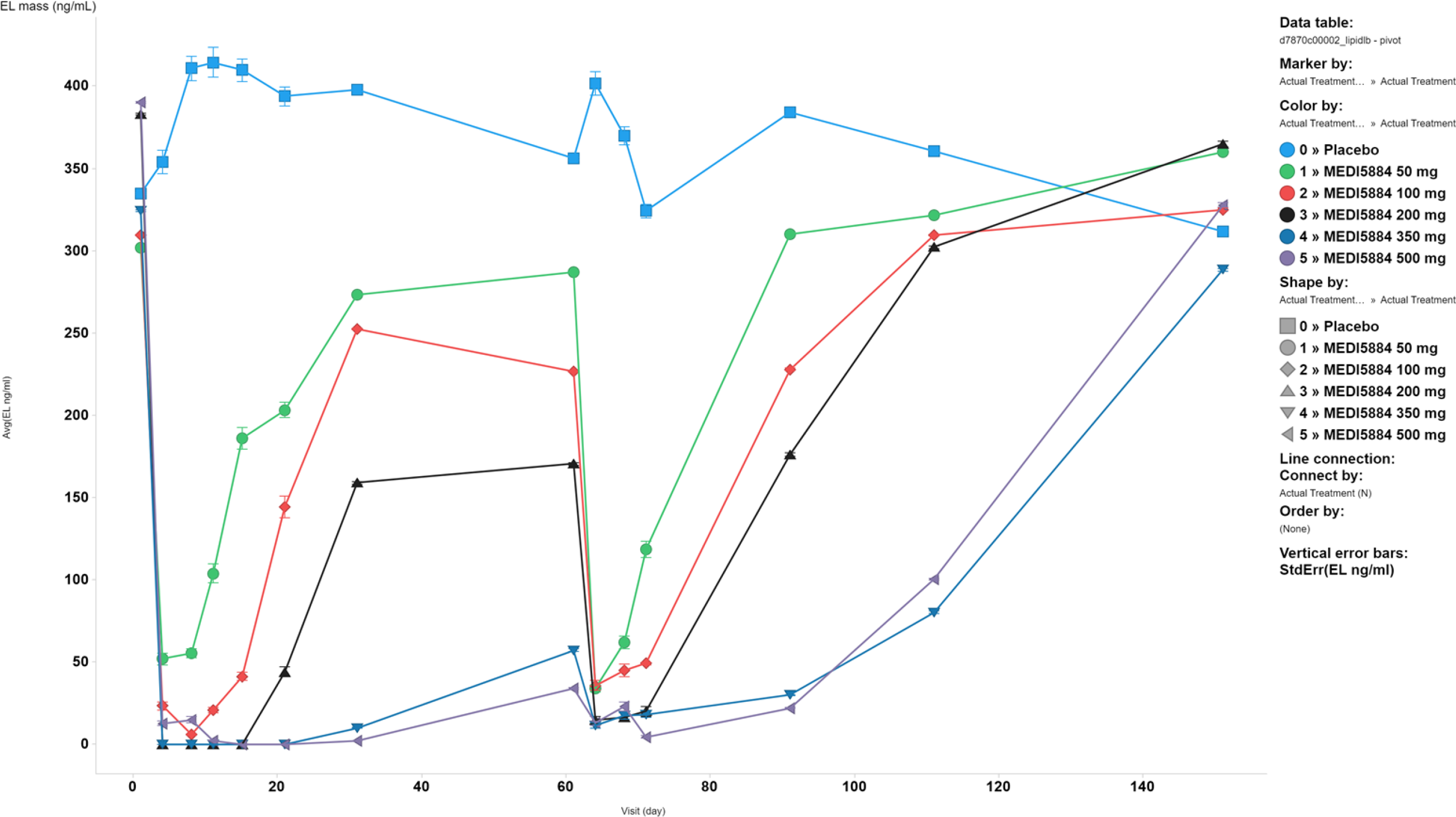
Dose-dependent suppression of EL levels (ng/mL) in CAD patients participating in study NCT03351738 upon administration of 3 monthly SC doses of MEDI5884. Avg = average; error bars represent SEM = standard error of the mean.

